# Oral administration of a single anti-CfaE nanobody provides broadly cross-protective immunity against major pathogenic Enterotoxigenic *Escherichia coli* strains

**DOI:** 10.1101/2020.06.16.155465

**Authors:** Alla Amcheslavsky, Aaron Wallace, Monir Ejemel, Qi Li, Conor McMahon, Matteo Stoppato, Serena Giuntini, Zachary A. Schiller, Jessica Pondish, Jacqueline R. Toomey, Ryan Schneider, Jordan Meisinger, Raimond Heukers, Andrew C. Kruse, Elieen M. Barry, Brian Pierce, Mark S. Klempner, Lisa A. Cavacini, Yang Wang

## Abstract

Enterotoxigenic *Escherichia coli* (ETEC) is estimated to cause approximately 380,000 deaths annually during sporadic or epidemic outbreaks worldwide. There is currently no vaccine licensed to prevent ETEC. Development of prophylaxis against ETEC is challenging due to the vast heterogeneity of the ETEC strains. The discovery of nanobodies has emerged as a successful new biologics in treating mucosal infectious disease as nanobodies can recognize conserved epitopes on hypervariable pathogens. In this study, we performed large screens using immunized llamas and a naïve nanobody yeast display library against adhesins of colonization factors. Cross-protective nanobodies were selected with *in vitro* activities inhibiting mannose-resistant hemagglutination (MRHA) against all eleven major pathogenic ETEC strains. Oral administration of nanobodies led to significant reduction of bacterial colonization in animals challenged with multiple ETEC strains. Structural analysis revealed novel conserved epitopes as critical structural features for pan-ETEC vaccine design.

Two of the lead nanobodies, 2R215 and 1D7, were further engineered as trimer or fused with human IgA Fc-fragments as fusionbodies. Oral administration of the trimers or fusionbodies protected mice from infection at a much lower dose compared to the monomeric format. Importantly, fusionbodies prevented infection as a pre-treatment when administrated 2 hours before ETEC challenge to the animals. Together, our study provides the first proof of concept that oral administration of a single nanobody could confer broad protection against major pathogenic ETEC strains. Technological advances in large*-*scale manufacturing of biological proteins in plants and microorganisms will make nanobody-based immunotherapy a potent and cost-effective prophylaxis or treatment for ETEC.

## Introduction

Enterotoxigenic *Escherichia coli* (ETEC) is one of the most common causes of diarrheal illness in children under 5 years of age, adults in the developing world and in travelers to endemic areas (1–3). ETEC infections are characterized by diarrhea, vomiting, stomach cramps, and in some cases mild fever. An estimated 10 million cases per year occur among travelers and military personnel deployed in endemic regions (3).When adult travelers develop ETEC diarrhea, a short course of antibiotics can reduce the duration and volume of diarrhea. However, ETEC strains are becoming increasingly resistant to antibiotics.

ETEC is a non-invasive pathogen that mediates small intestine adherence through bacterial surface structures known as colonization factors (CFs). Once bound to the small intestine, the bacteria produce toxins causing a net flow of water from enterocytes, leading to watery diarrhea. Development of an effective vaccine against ETEC bacterial attachment and colonization has long been considered as a promising approach against ETEC diarrhea. CFA/I is one of the most prevalent CFs and is composed of CfaE, the tip minor adhesin subunit, and a homopolymeric structural subunit, CfaB. In human clinical trials, oral administration of anti-CfaE bovine IgG provided protection against virulent ETEC challenge in over 60% of the test group, suggesting that an adhesin-based vaccine could be effective to elicit endogenous production of protective antibodies (4). Recent studies in our laboratories demonstrated that when administered orally to mice and non-human primates, anti-CfaE human monoclonal antibodies inhibited bacterial colonization in the small intestine and protected animals from diarrhea associated with ETEC infection (5).

ETEC strains are antigenically diverse with over 25 types of CFs having been identified (6–9). CFA/I represent the archetype of class 5 fimbriae, the largest class of human specific CFs that causes a majority of moderate to severe ETEC diarrheal cases (7, 9, 10). Class 5 fimbriae includes three sub-classes, 5a (CFA/I, CS4, CS14), 5b (CS1, CS17, CS19, PCF071), and 5c (CS2). Other non-class 5 fimbriae have also been implicated in endemic and traveler diarrheal diseases including helical CS5, fibrillary CS3, CS21 and non-fimbrial CS6 adhesins (11). As such, development of a broadly protective vaccine against all major pathogenic ETEC strains remains one of the most important challenges in vaccine design (12, 13). In theory, a cross-protective vaccine could be generated with a combination of intact CFs (14). However, such strategies could lead to unexpected instability or safety concerns and high costs for manufacturing production. Despite advances in anti-ETEC vaccine research, there remains no licensed vaccine against ETEC.

The discovery of camelid heavy-chain variable domains (VHHs or nanobodies) has produced a new application for antibody-derived biologics (15). Nanobodies consist of a highly stable and soluble single antigen-binding variable region (15 kDa), since they are derived from heavy chain only antibodies with no associated light chains (16). Unlike conventional antibodies, nanobodies can access distinct antigenic sites (e.g., enzyme active sites, recessed regions of viral glycoproteins) due to smaller paratope diameters and a longer complementarity-determining region 3 (CDR3) to form finger-like extensions (15, 17). Based on their unique characteristics, nanobodies can recognize conserved epitopes on hypervariable pathogens, such as HIV, poliovirus, norovirus, and coronavirus (18–22). Consequently, nanobodies are being studied in clinical trials for the treatment of a wide range of diseases by both systemic and oral administration due to their excellent solubility and thermostability (23–25).

In this study, we identified a panel of nanobodies that showed broad protective activities against eleven major types of disease-causing strains of ETEC. Two of the lead nanobodies, 2R215 and 1D7, were engineered as trimer or fused with human IgA Fc-fragments to generate fusionbodies (VHH-IgA Fc) to further improve the potency. Oral administration of the trimers and fusionbodies prevented and protected mice from infection at a much lower dose compared to monomeric nanobodies.

## Results

### Identification of anti-ETEC nanobodies from immunized llama and yeast display library

To generate nanobodies with broad cross-reactivity against ETEC adhesins, two male llama were subcutaneously immunized with N-terminal fragments of eight class 5 ETEC adhesins (**Figure 1A**). Serum response to adhesins was measured by ELISA and PBMC were isolated from llama to generate phage-displayed libraries of size around 10^8^ as previously described (26). Libraries were initially used for panning on immobilized colonization factor antigens CS1 and CS2, because of their relatively low sequence homology. Output from those pannings were used for a subsequent panning round on CfaE. Periplasmic extracts were prepared and tested for binding to all 8 adhesin antigens in ELISA. Cross-reactive clones were sent for sequence determination and further characterization. From these selections, 26 candidate nanobodies were selected.

**Figure 1.**
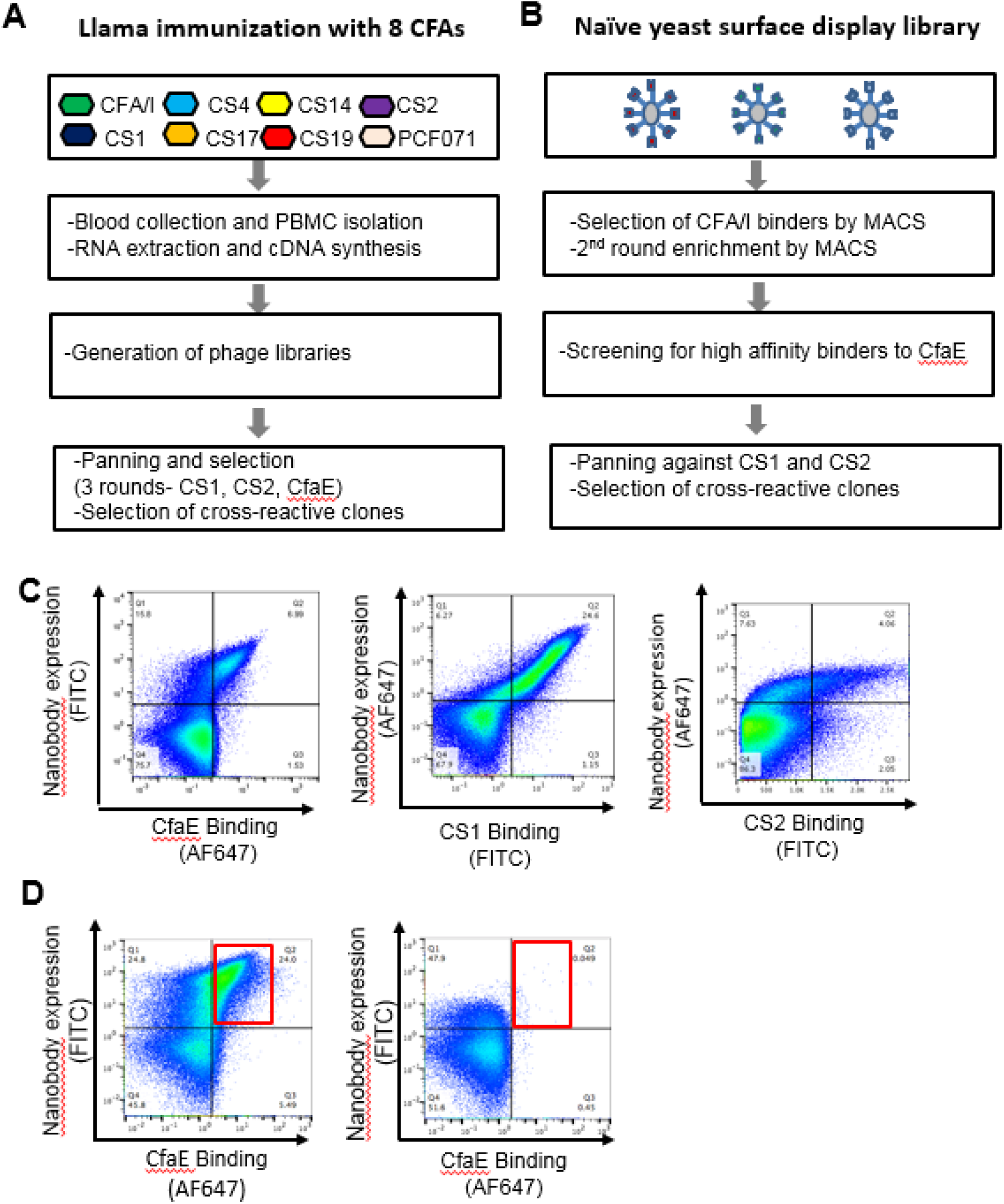
Identification of anti-ETEC nanobodies from immunized llama and yeast display library. (A) Flowchart of llama immunization and panning. Two male llama were subcutaneously immunized with N-terminal fragments of eight class 5 ETEC adhesins. PBMC were isolated from llama to generate phage-displayed libraries of approximately 10^8^ in size. Libraries were initially panned on immobilized colonization factor antigens CS1 and CS2. Output from those pannings was used for a subsequent panning round on CfaE. Cross-reactive clones were sent for sequence determination and further characterization. (B) Flowchart of selection process of synthetic nanobodies. Binders to CfaE from the naïve yeast-displayed library were enriched over two rounds of selection by performing magnetic-activated cell sorting (MACS) on yeast that were stained with purified and labeled CfaE. High affinity binders were selected during fluorescence-activated cell sorting (FACS) and single cell clones were screened for binding with CS1 and CS2. Cross-reactive clones were sent for sequence determination and further characterization. (C) Flow analysis demonstrating example of binding of yeast displaying nanobody 2R23 to CfaE, CS1 and CS2 proteins. (D) Flow analysis demonstrating functional human monoclonal antibody 68-61 competing for binding to CfaE with yeast library derived nanobody 2R215 on yeast.

In parallel, we used a recently developed library of synthetic nanobodies displayed on the surface of *Saccharomyces cerevisiae* (27). Two rounds of magnetic activated cell sorting (MACS) were performed using FITC and AlexaFluor647-labeled N-terminal fragments of CfaE to identify CfaE binding clones (**Figure 1B**). High-affinity binders were enriched by fluorescent activated cell sorting (FACS) with decreasing concentration of FITC labeled CfaE as previously described (27, 28). Approximately 300 yeast clones were recovered from the yeast library screen. Binding of selected clones to antigen was verified by yeast surface staining against AlexaFluor647 labeled CfaE using flow cytometry. The clones showed a range of binding activities to CfaE **(Supplemental figure 1)**. All binders were subjected to additional rounds of screening for cross-reactivity against two other class 5 adhesins, CS1 and CS2 (**Figure 1C**). A total of 30 nanobodies with cross-reactivity and unique sequences were selected for further characterization. In addition, a competitive FACS screen was performed against a potent anti-CfaE full length antibody, 68-61, to identify nanobodies specific to the receptor binding region of CfaE (29). One clone, 2R215, was found to recognize an overlapping epitope with 68-61 (**Figure 1D, Supplemental figure 2**).

### Selected anti-CfaE nanobodies show cross-reactivity against all class 5 colonization factors

To test the breadth of cross-reactivity, selected llama- and yeast-derived nanobodies were sequenced, cloned in pET-26b vector, expressed in *E.coli*, and examined by ELISA for binding to eight class 5 adhesins. Two yeast-derived clones (2R215 and 2R23) and two llama-derived clones (1D7 and 1H4) were selected based on their broad reactivity against multiple adhesins (**Figure 2A-D**). The four lead nanobodies were then tested for inhibition of mannose resistant hemagglutination (MRHA) against an ETEC strain expressing CFA/I (H10407), the most prevalent class 5 adhesin. All nanobodies showed maximal inhibitory concentration (IC100) in the micromolar concentration range (1.9 to 3.7μM) (**Figure 2E**). We next examined if these nanobodies had activity against ETEC strains expressing 7 other class 5 adhesins in MRHA assay. Consistent with ELISA binding data, the four nanobodies were cross-protective against all eight strains with an IC100 activity ranging from 0.4125 to 13.2 μM (**Table 1**).We next examined the MRHA activity of nanobodies against ETEC strains expressing other pathogenic fimbrial and non-fimbrial adhesins including helical CS5, fibrillary CS3, and non-fimbrial CS6 adhesins, as well as another common diarrhea causing adhesin CS21. Surprisingly, four nanobodies showed broad protection against all tested strains with an IC100 ranging from 0.2063 to 6.6 μM (**Table 1**).

**Figure 2.**
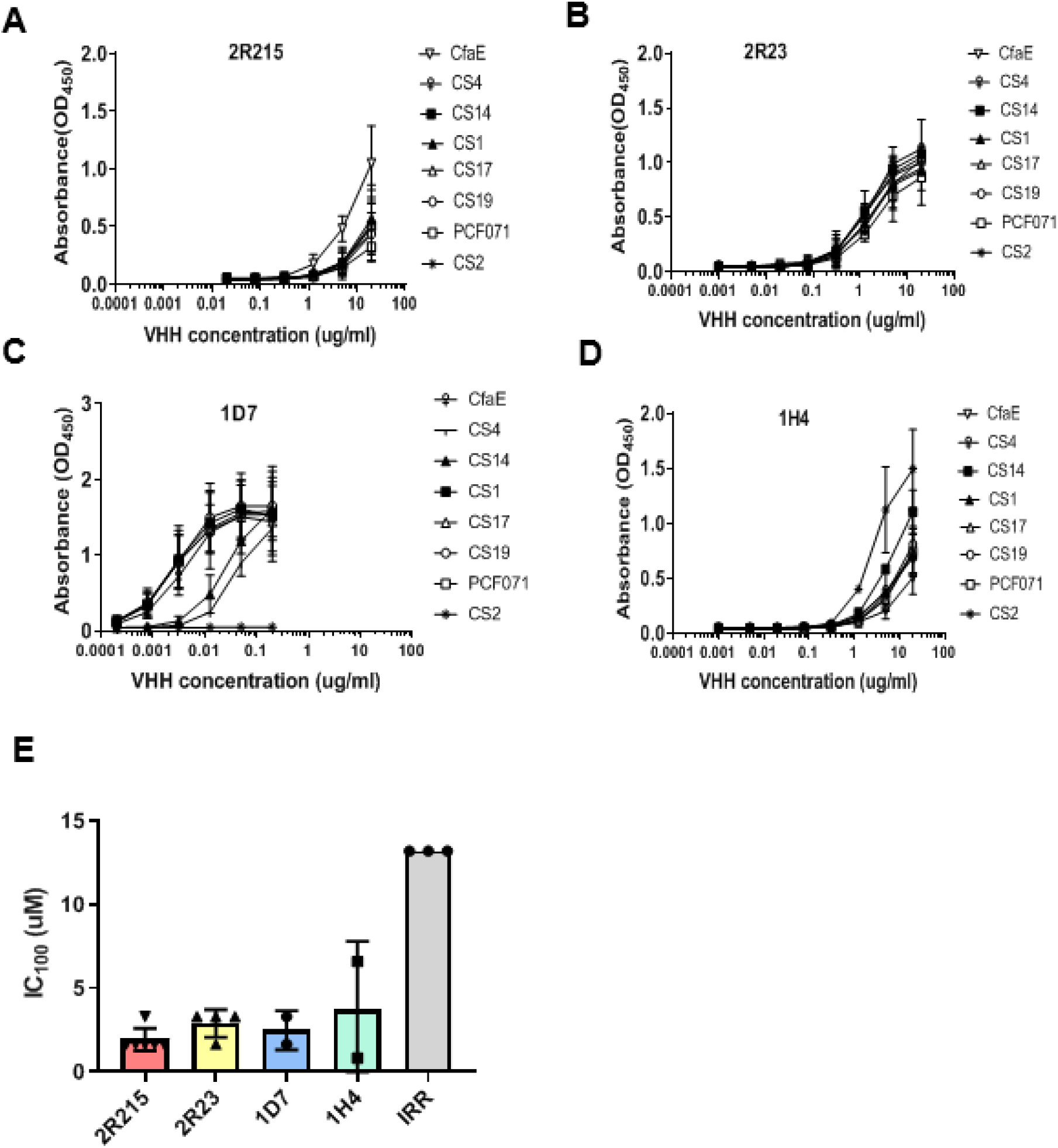
Selected anti-CfaE nanobodies show cross-reactivity against class 5 colonization factors. (A-D) Two yeast-derived clones (2R215 and 2R23) and two llama-derived clones (1D7 and 1H4) were examined by ELISA for binding to eight class 5 adhesins. These clones showed broad reactivity against multiple adhesins. (E)The four lead nanobodies were then tested for inhibition of mannose resistant hemagglutination (MRHA) against an ETEC strain expressing CFA/I (H10407). All nanobodies showed maximal inhibitory concentration (IC100) in the micromolar concentration range (1.9 to 3.7μM).

**Table 1.** Lead nanobodies are broadly cross-functional against multiple adhesins. (A) Four nanobodies were examined for activity in MRHA assay against eight strains representing class5 of ETEC antigens and other pathogenic fimbrial and non-fimbrial adhesins CS5, CS3, CS6 and CS21. Nanobodies showed broad protection against all tested strains with an IC_100_ ranging from 0.31 to 16.67 μM.

|  | Rigid rods |  |  |  |  |  |  | Helical | Fibrillar | Non fimbrial | Bundle-forming |
| --- | --- | --- | --- | --- | --- | --- | --- | --- | --- | --- | --- |
|  | 5a |  |  | 5b |  |  | 5c | CS3 | CS5 | CS6 | CS21 |
|  | CFA/I | CS4 | CS14 | CS1 | CS17 | CS19 | CS2 |  |  |  |  |
| VHH |  |  |  |  |  |  |  |  |  |  |  |
| 2R215<br>IC<sub>100</sub> (uM) | 4.17±1 | 5.5±1.37 | 13.3±0 | 13.3±0 | 1.31±1.02 | 9.44±3.96 | 8.75±2.84 | 5±1.68 | 1.04±0.6 | 3.89±1.82 | 1.25±0.59 |
| 2R23<br>IC<sub>100</sub> (uM) | 10±0 | 2.22±0.56 | 4.17±2.53 | 13.3±0 | 0.76±0.46 | 9.44±3.96 | 2.60±1.39 | 10±3.37 | 1.11±0.28 | 2.92±1.38 | 0.83±0.42 |
| 1D7<br>IC<sub>100</sub> (uM) | 16.67±3.35 | 7.29±3.49 | 6.65±0 | 5±1.68 | 0.31±0.10 | 5±1.7 | 0.94±0.63 | 3.6±1.68 | 0.97±0.74 | 3.61±1.72 | 0.97±0.37 |
| 1H4<br>IC<sub>100</sub> (uM) | 7.5±2.1 | 1.35±0.31 | 6.65±0 | 5.55±1.48 | 0.76±0.46 | 7.2±3.44 | 2.7±0.62 | 5±1.68 | 0.52±0.32 | 4.17±2.52 | 1.25±0.42 |

### Epitope mapping of lead anti-ETEC VHHs

To elucidate the interaction between anti-ETEC nanobody and adhesin at the molecular level, we performed *in silico* modeling of four nanobody/adhesin complexes by docking simulations and alanine scanning. Computational binding models were built based on the previously resolved crystal structure of CfaE (PDB ID: 2HBO), and homology modeling with the structure of 6H16 nanobody (PDB ID: 6H16) for llama-derived 1D7 and 1H4, and 5VNV nanobody (PDB ID: 5VNV) for yeast-derived 2R215 and 2R23 using BioLuminate (*Schrodinger*) (**Figure 3A-D**).

**Figure 3.**
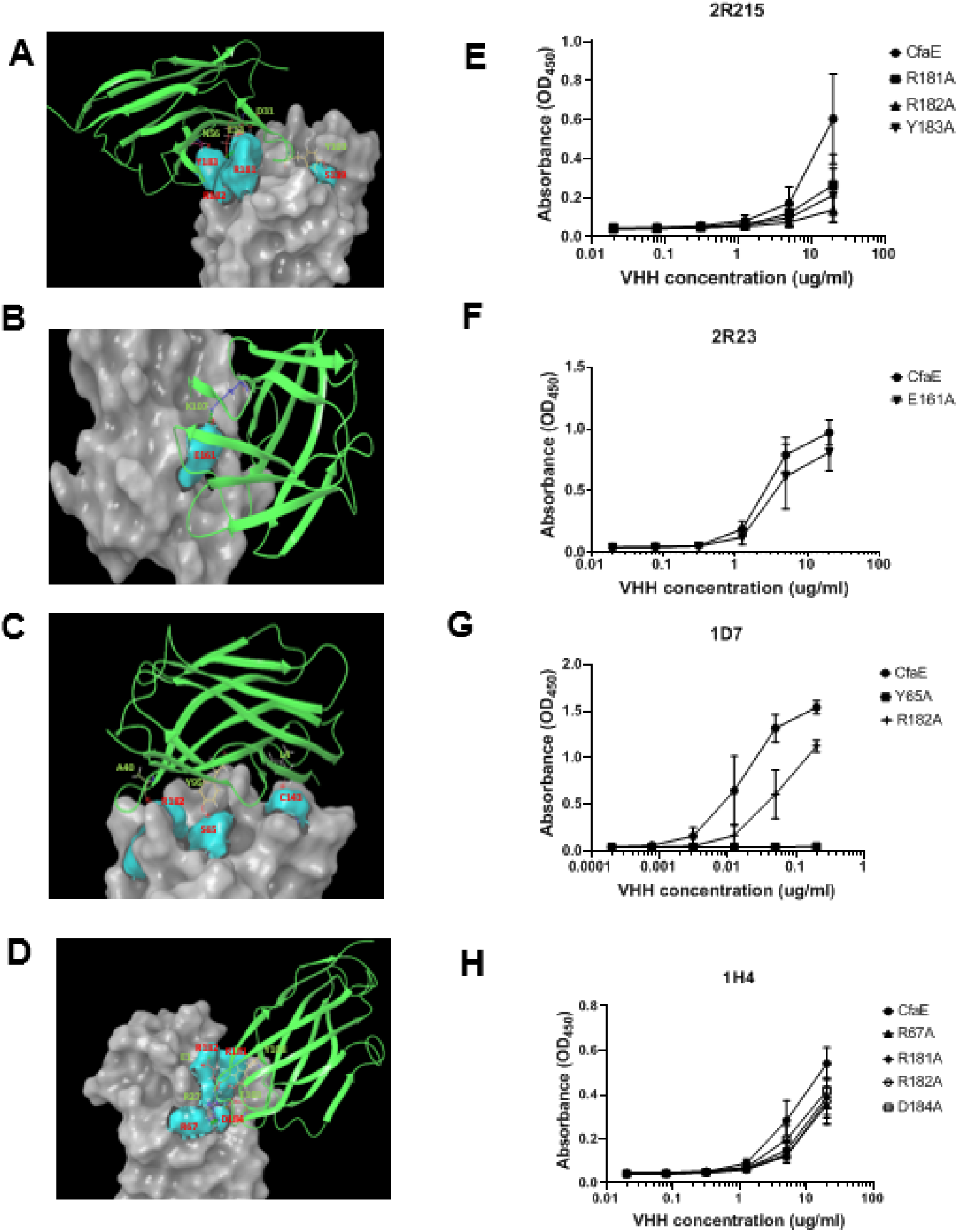
Epitope mapping of lead anti-ETEC VHHs. (A-D) Computational binding models were built based on the previously resolved crystal structure of CfaE (PDB ID: 2HBO), and homology modeling with the structure of 6H16 nanobody (PDB ID: 6H16) for llama-derived 1D7 and 1H4, and 5VNV nanobody (PDB ID: 5VNV) for yeast-derived 2R215 and 2R23 using BioLuminate (*Schrodinger*) (E-F) Individual CfaE residues were selected for mutagenesis to alanine and tested by ELISA for nanobody binding. Mutating Y65, R67, E161, D184, R181, R182, and Y183 to alanine affected the binding of nanobodies to CfaE suggesting that these residues are involved in the binding interactions.

Based on docking results, individual CfaE residues within or adjacent to the putative receptor binding pocket formed by three arginine residues (R67, R181 and R182) were selected for mutagenesis to alanines and tested by ELISA for nanobody binding (**Figure 3E-H**). ELISA results showed that mutating Y65, R67, E161, D184, R181, R182, and Y183 to alanine affected the binding of nanobodies to CfaE suggesting that these residues are involved in the binding interactions. This is in line with findings by Baker who demonstrated R181 of CfaE as crucial for ETEC binding in a human in vitro organ culture (30). In addition to the residues near the putative receptor domain, we identified several other highly conserved residues such as S65, L64, Y58, L110, Y156 and C143 that are required for interaction between nanobodies and adhesin. Interestingly, sequence alignment revealed that these residues are highly conserved among at least eight class 5 adhesins (31), which is consistent with our observation that these nanobody are broadly cross protective against multiple ETEC strains.

### Anti-CfaE VHH prevent ETEC colonization in the small intestine of a mouse model

The lead nanobodies were further evaluated in a mouse model of ETEC colonization as described previously (29) (**Figure 4A**). Groups of five DBA/2 mice were given a mixture of bacteria and anti-CfaE nanobodies by oral gavage. All nanobodies were tested at 100 mg/kg, except for 2R23, which was tested at 50mg/kg due to the low production yield. Twenty four hours after challenge, the mice were euthanized and colony forming units (CFUs) in the small intestine were counted to determine the level of bacterial colonization. The efficacy of the anti-CfaE nanobodies was assessed by determining whether nanobodies could prevent adhesion/colonization of bacteria to the small intestine compared to an irrelevant nanobody control. In the anti-CfaE nanobody treatment groups, there was a log reduction of CFUs by 2.9 for 2R215, 2.3 for 1H4, 1.7 for 1D7 and 1.5 for 2R23 compared to the irrelevant nanobody control group (*P* <0.01 for 2R215 and *P*<0.05 for 2R23,1D7 and 1H4). These results indicate that all four nanobodies showed significant activity in preventing colonization by H10407 strain at a dose of 50 or 100 mg/kg (**Figure 4B**).

**Figure 4.**
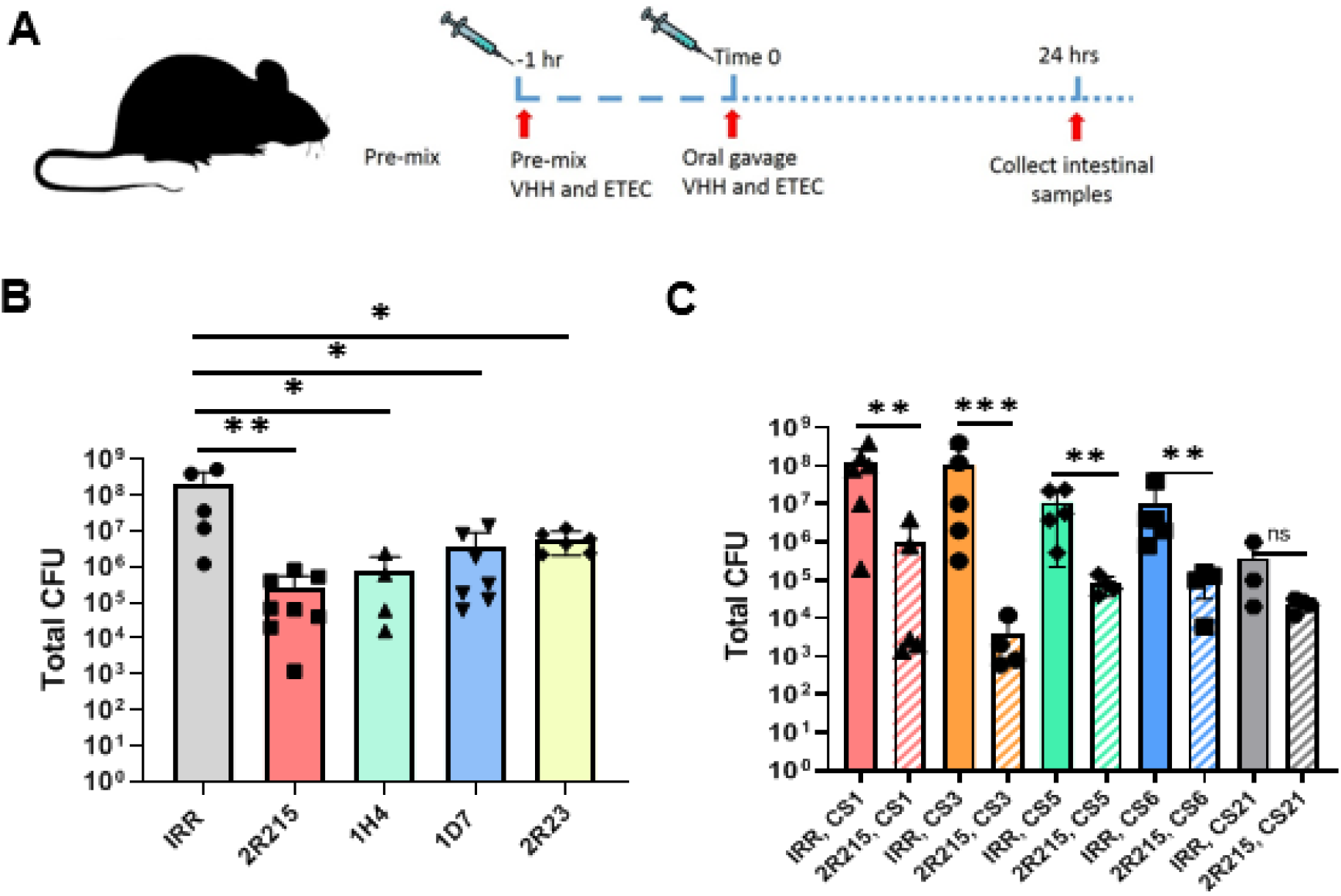
Anti-CfaE VHH prevent ETEC colonization in the small intestine of a mouse model. (A) Cartoon representation of mouse colonization experiment. VHH were pre-incubated with 10^7^ colony forming units (CFU) of ETEC strains 1 hour prior to intragastric challenge. Animals are euthanized 24 hours after challenge, and bacterial colonies in the small intestine are counted. (B) DBA/2 mice were challenged intragastrically with 10^7^ CFU of H10407 pre-incubated with 100mg/kg of VHH. For 2R23 50mg/kg of VHH was used due to lower production yields of this nanobody. At least 5 animals were tested for each condition. Results for all VHH were significantly different from those for the irrelevant VHH (*P* <0.01 for 2R215 and *P*<0.05 for 2R23, 1D7 and 1H4). (C) DBA/2 mice were challenged intragastrically with 10^7^ CFU of strains expressing H10407 adhesins CS1, CS3, CS5, CS6, and CS21 pre-incubated with 100mg/ml of 2R215. Treatment with 2R215 resulted in significant reduction of CFUs against ETEC strains expressing CS1 (2.1 log, *P*<0.01), CS3 (4.4 log, *P*<0.001), CS5 (2.2 log, *P*<0.01) and CS6 (2 log, *P*<0.01), but not the strain expressing CS21 (1.2 log, *P*>0.05)

The lead nanobody, 2R215, was then tested against pathogenic ETEC strains expressing other adhesins that are commonly isolated from patients with ETEC-associated diarrheal diseases, including CS1, CS3, CS5, CS6, and CS21 (9, 11). 2R215 demonstrated broad protection against all tested strains. Compared to the irrelevant control, treatment with 2R215 at 100mg/kg resulted in significant reduction of CFUs against ETEC strains expressing CS1 (2.1 log, *P*<0.01), CS3 (4.4 log, *P*<0.001), CS5 (2.2 log, *P*<0.01) and CS6 (2 log, *P*<0.01), but not the strain expressing CS21 (1.2 log, *P*>0.05) (**Figure 4C**).

### Multimerization enhances the potency of 2R215 in vitro and in vivo

Multimerization of single domain nanobodies can increase their stability and potency (26, 32). To improve the potency of identified cross protective nanobodies, 2R215 and 1D7 were engineered as a dimer or trimer molecules using 3x or 6x (G4S) linkers in tandem N to C terminus orientation (**Figure 5A**). Multimerized nanobodies were expressed in bacteria and purified using fused His tags (**Figure 5B**). Both dimeric and trimeric 2R215 showed increased potency in Caco2 cell adhesion assays. The IC50 values are 10.213 μM for monomeric 2R215, 5.09 μM for dimeric 2R215 and 1.06 μM for trimeric form of 2R215. (**Figure 5C**). Activity in the MRHA assay was also improved with an IC100 of 6.7 μM for monomeric, 1.25 for dimeric and 2.5 for the trimeric form of 2R215 (*P*<0.01 and *P*<0.05 for dimeric and trimeric forms respectively) (**Figure 5D**).

**Figure 5.**
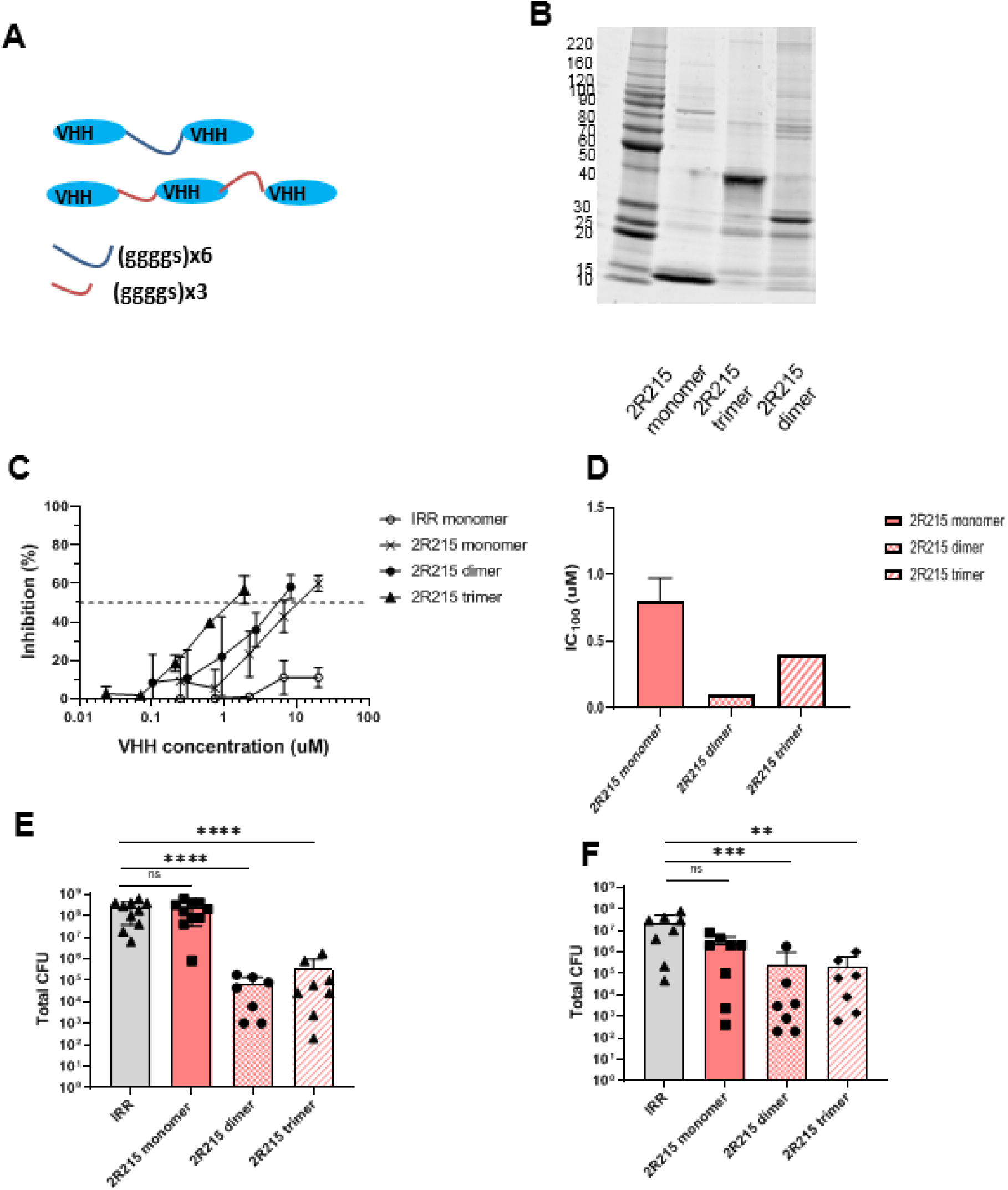
Multimerization enhances the potency of 2R215 in vitro and in vivo. (A) Cartoon representation of multimerization strategy. VHH 2R215 was engineered as a dimer or trimer molecules using 3x or 6x (G4S) linkers in tandem N to C terminus orientation. (B) Multimerized nanobodies were expressed in bacteria and purified using fused His tags. Purified multimer sizes were verified by polyacrylamide gel electrophoresis. (C) Dimeric and trimeric 2R215 show increased potency in Caco2 cell adhesion assays. The IC50 values are 10.213 μM for monomeric 2R215, 5.09 μM for dimeric 2R215 and 1.06 μM for trimeric form of 2R215. (D) Activity in the MRHA assay is improved with an IC100 of 6.7 μM for monomeric, 1.25 for dimeric and 2.5 for the trimeric form of 2R215 (*P*<0.01 and *P*<0.05 for dimeric and trimeric forms respectively). (E) Mice inoculated with ETEC mixed with 10 mg/kg of dimeric or trimeric forms 2R215 showed a log reduction of 3.5 and 2.8 in colonization (*P*<0.0001) compared to the monomeric 2R215 treated group at the same dose (F) The dimeric and trimeric forms of 2R215 retained their activity when administered 2 hours prior to bacteria challenge as compared to the monomeric form. Pre-treatment with dimeric and trimeric 2R25 reduced total CFUs by 1.9 log (*P*<0.001) and 2 log (*P*<0.01) respectively.

The dimeric and trimeric forms of 2R215 were further characterized in a mouse colonization model. Mice inoculated with ETEC mixed with 10 mg/kg of dimeric or trimeric forms 2R215 showed a log reduction of 3.5 and 2.8 in colonization (*P*<0.0001) compared to the monomeric 2R215 treated group at the same dose (**Figure 5E**). Moreover, the dimeric and trimeric forms of 2R215 retained their activity when administered 2 hours prior to bacteria challenge as compared to the monomeric form. Pre-treatment with dimeric and trimeric 2R25 reduced total CFUs by 1.9 log (*P*<0.001) and 2 log (*P*<0.01) respectively (**Figure 5F).** These results indicated that multimerization of monomeric nanbodies may result in improved intragastric stability and protective potency.

### IgA Fc fusion enhances in vivo potency of 2R215 and 1D7

To gain effector function and improve mucosal stability, nanobodies could be engineered with the IgA Fc domain as IgA Fc fusionbodies (VHH-IgA) for oral administration (33). To engineer VHH-IgA Fc fusionbodies, 2R215 and 1D7 were grafted on to the Fc domain of IgA1 and IgA2 (VHH-IgA1 and VHH-IgA2) at the hinge regions as bivalent VHH-Fc fusionbodies (34) (**Figure 6A**). 2R215 and 1D7 VHH-IgA1 and VHH-IgA2 fusionbodies were expressed in Expi293 cells and the purified fusion protein sizes were verified by polyacrylamide gel electrophoresis (**Figure 6B)**. The binding of fusionbodies with adhesin was maintained with increased binding activities as compared to monomeric nanobody 2R215 and 1D7 in ELISA **(Figure 6C, D)**.

**Figure 6.**
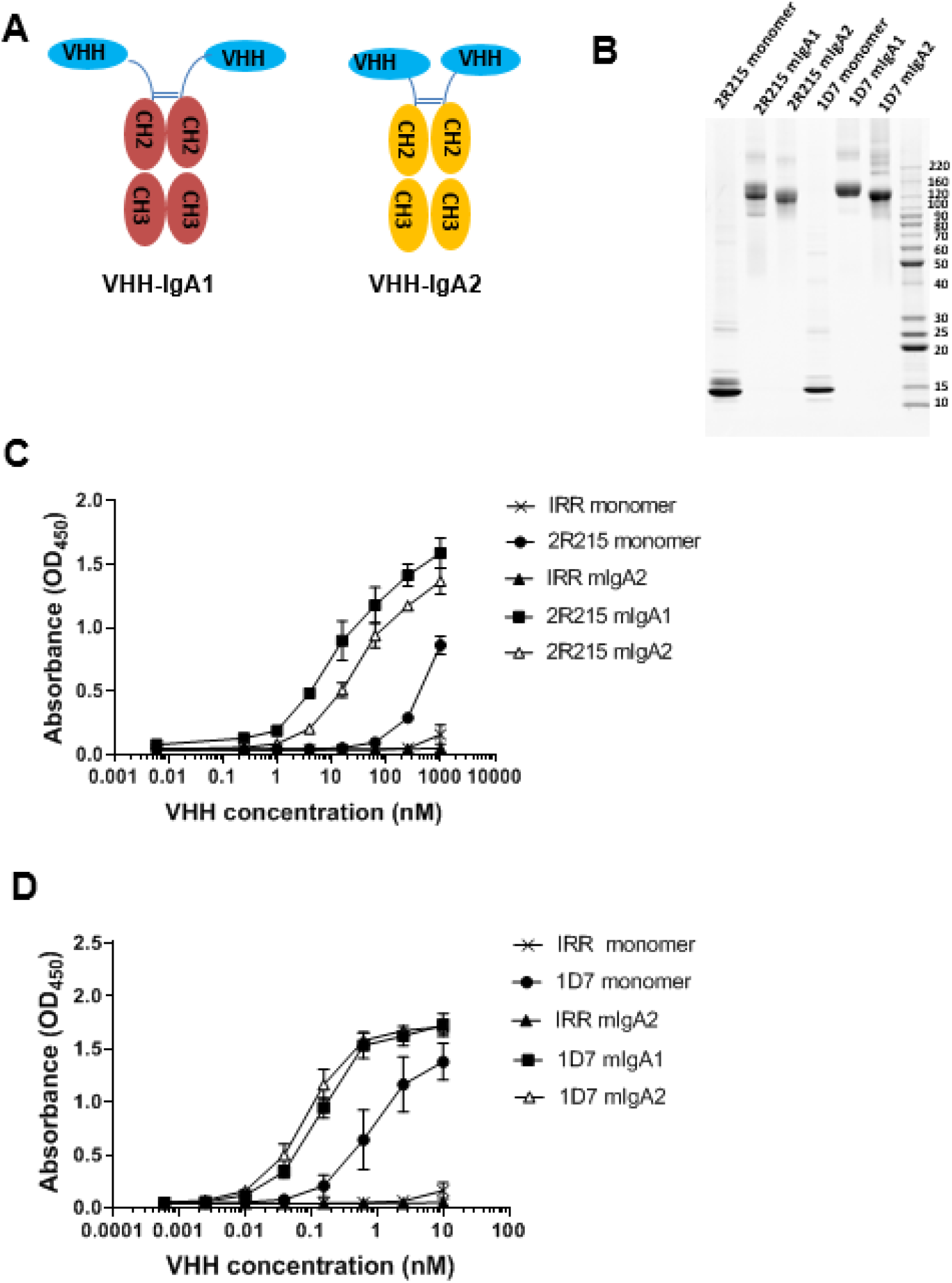
IgA Fc fusion enhances in vitro potency of 2R215 and 1D7. (A) Cartoon representation of IgA fusion strategy. To engineer VHH-IgA Fc fusionbodies, 2R215 and 1D7 were grafted on to the Fc domain of IgA1 and IgA2 (VHH-IgA1 and VHH-IgA2) at the hinge regions as bivalent VHH-Fc fusionbodies. (B) 2R215 and 1D7 VHH-IgA1 and VHH-IgA2 fusionbodies were expressed in Expi293 cells and the purified fusion protein sizes were verified by polyacrylamide gel electrophoresis. (C, D) ELISA showing the binding of fusionbodies 2R215 and 1D7 with CfaE. The binding of fusionbodies with adhesin was maintained with increased binding activities as compared to monomeric nanobodies.

2R215 and 1D7 fusionbodies were then tested in mouse colonization assay administered with ETEC strain H10407. While monomeric 2R15 and 1D7 inhibited colonization at 100 mg/kg **(Figure 4B)**, the IgA fusionbodies showed stronger inhibitory activity at a much lower dose of 10 mg/kg (**Figure 7A-D**). Next, the most potent fusionbodies, 1D7 VHH-IgA1 (3.5 log reduction in CFU, *P*<0.001) and 2R215 VHH-IgA2 (5 log reduction in CFU, *P*<0.001), were examined in a pre-treatment model. Animals were orally inoculated with fusionbodies 1 hour prior to ETEC challenge. Fusionbody treatment resulted in reduction of CFUs by 1.2 log for 2R215 VHH-IgA2 (*P*<0.01) and 2.7 log for 1D7 VHH-IgA1 (*P*<0.0001) (**Figure 7C, D**). Furthermore, the 1D7 VHH-IgA1 fusionbody retained activity when the pre-treatment time was extended to 2 hours. A 1.7 log reduction in total CFUs was observed in 1D7 VHH-IgA1 treated animals as compared to the group treated with irrelevant control (*P*<0.0001) (**Figure 7E**). We also compared the activity of the 1D7 VHH-IgA1 fusionbody to Travelan, a commercial hyperimmune bovine colostrum (HBS) product used for prevention of ETEC-induced diarrhea. While Travelan showed activity in 1 hr pre-treatment when used at 40mg/kg **(Supplemental Figure 3)**, it did not inhibit colonization after 2 hours of pre-treatment **(Figure 7E)**.

**Figure 7.**
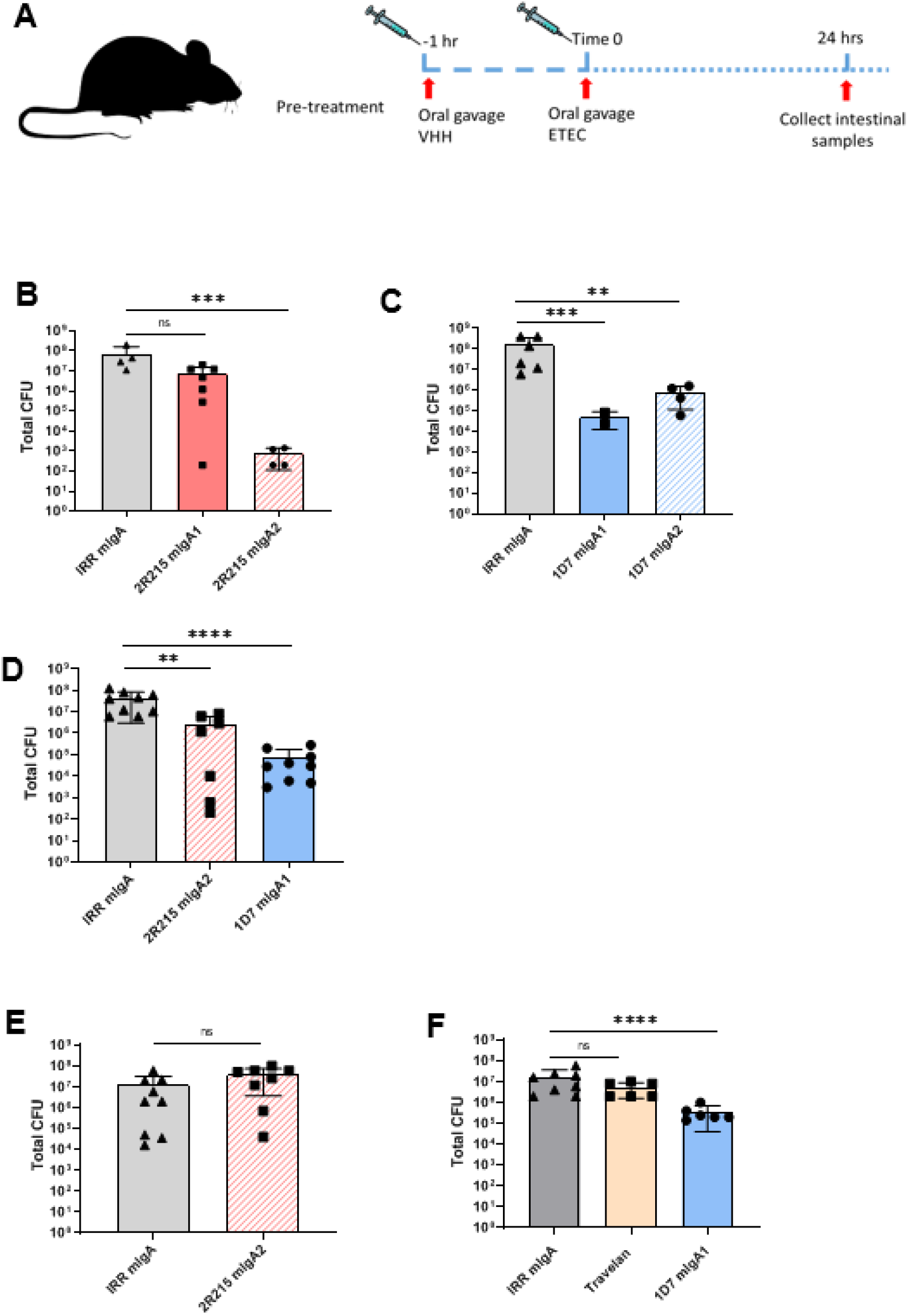
IgA Fc fusion enhances in vivo potency of 2R215 and 1D7. (A) Cartoon representation of pre-treatment protocol of VHH administration. VHH are administered one or two hours prior to administration of 10^7^ CFU of H10407. Animals are euthanized 24 hours after challenge, and bacterial colonies in the small intestine are counted. (B, C) DBA/2 mice were challenged intragastrically with 10^7^ CFU of H10407 pre-incubated with 10 mg/kg of VHH. Treatment with 1D7 VHH-IgA1 resulted in 3.5 log reduction in CFU, *P*<0.001) and 2R215 VHH-IgA2 in 5 log reduction in CFU (*P*<0.001). (D) Fusionbodies 2R215 and 1D7 were intragastrically administered one hour prior to administration of 10^7^ CFU of H10407. Fusionbody pre-treatment resulted in reduction of CFUs by 1.2 log for 2R215 VHH-IgA2 (*P*<0.01) and 2.7 log for 1D7 VHH-IgA1 (*P*<0.0001). (E, F) Fusionbodies 2R215 and 1D7 were intragastrically administered two hours prior to administration of 10^7^ CFU of H10407. A 1.7 log reduction in total CFUs was observed in 1D7 VHH-IgA1 treated animals as compared to the group treated with irrelevant control (*P*<0.0001).

## Discussion

ETEC strains are highly antigenically diverse, expressing over 25 types of adhesion factors. It has been very challenging to generate a pan-ETEC vaccine with broad coverage against diverse strains that cause diarrheal disease. In the present study, we used a llama immunization and a synthetic yeast library screen to identify and characterize a panel of nanobodies that recognize highly conserved epitopes on hypervariable adhesins. Among the 50 isolated nanobodies, four lead nanobodies showed broad coverage against 11 pathogenic ETEC strains and protected animals from infection.

We previously described a positively charged binding pocket at the N-terminal domain of CfaE important for ETEC interaction with the negatively charged sialic acid on the host epithelial cells (29). This binding pocket is formed by three arginine residues, R67, R181, and R182, and a cluster of surrounding residues, which are all highly conserved among the class 5 fimbrial adhesins. Mutations of the arginine residues result in a complete loss of binding activity of ETEC to host cells (30). Yeast library derived nanobody, 2R215, was found to directly bind to R181 in the putative receptor binding domain. 1D7 and 1H4 derived from the immunized llama were also found to bind directly to R182 (part of the putative receptor binding domain). 2R215, 1D7, and 1H4 recognized three other residues, Y183, Y65, and C143, within or near the receptor binding domain. Based on structural analysis, three highly conserved residues in class 5 adhesins, P46, Y58, and Y156, were shared epitopes for all four cross-protective nanobodies. Taken together, our structural finding identified novel conserved domains on various ETEC CFs as critical features of an effective pan-ETEC vaccine.

Multimerization and IgA Fc fusion of nanobodies has emerged as an effective way to increase potency and mucosal half-life (26). Recent studies have shown the feasibility of producing various formats of nanobodies in plants and microorganisms. High levels of VHH-IgA fusionbodies were effectively produced in yeast and soybean and have been proven efficient for mucosal protection against ETEC in piglets (34, 35). Moreover, an anti-rotavirus VHH (ARP1) produced in yeast and rice was capable of protecting mouse pups against rotavirus-induced diarrhea (36, 37). Oral administration of ARP1 produced in yeast was safe and effective in reducing diarrhea in male infants with severe rotavirus-associated diarrhea (38).

In the absence of a pan-ETEC vaccine, our study provides the first proof of concept that oral administration of a nanobody, prior to or during infection, could confer broad protection against various ETEC strains. Compared to Travelan, a commercial HBC product used to prevent ETEC-induced diarrhea, 1D7 VHH-IgA1 fusionbody appears to be more stable in mouse colonization model when administered two hours prior to ETEC challenge and is effective at lower dose. Technological advances in large-scale manufacturing of biological proteins in plants and microorganisms will make nanobody-based immunotherapy a potent and cost-effective prophylaxis or treatment for ETEC.

## Materials and Methods

### Antigen cloning, expression, and purification

The nucleic acid sequences of N-terminal adhesin domains of CfaE and class 5 adhesins (GenBank M55661) were cloned into a pMAL-C5X vector (Addgene) in-frame with an MBP tag to express as periplasmic proteins with improved solubility (MBP–CfaE-N). The donor strand complement was included to ensure the overall protein expression and stability, as reported previously(39). All cloned constructs were transformed into SHuffleT7 Competent Escherichia coli (NEB), and expression was induced with 1 mM IPTG (isopropyl-D-thiogalactopyranoside). Bacteria were lysed, and proteins were purified with amylose resin (NEB) and eluted with 20 and 50 mM maltose (Sigma).

### Llama immunization

Immunizations of llamas and library constructions were performed as described previously (26). Briefly, two male llamas were immunized with antigen cocktails that included eight class 5 colonization factors (300ug each antigen, 2400ug total for each animal). The first cocktail consisted of CS2, CS4, CS14, CfaE and the second cocktail consisted of CS19, CS17, CS1, and PCF071. The two antigen cocktails were injected to each animal weekly for six weeks and 150 ml blood was collected for isolation of RNA from the peripheral blood lymphocytes after the last immunization on the 7^th^ week. Good immune response was observed in both animals against all antigens. RNA isolation and library constructions were performed by amplifying nanobody genes and ligating into a phagemid vector. Two phage display libraries were generated with a size of approximately 5×10^8^ transformed *E. coli* TG1 bacteria and a correct insert ration of ~100%. Selection of target-binding nanobodies was performed by phage-display selections against immobilized antigen proteins on Maxisorp plates (NUNC, Thermo Fisher Scientific). The initial screening was performed against CS1and CS2, followed by a second round of screening against the CfaE antigen. Phages were eluted by unspecific and competitive elutions. Eluted phage were used to infect exponentially growing *E. coli* TG1 that were plated on Luria broth (LB) agar plates containing 2% (w/v) glucose and 100 μg/ml ampicillin. In addition, eluted phages were serially diluted, infected in TG1 and 5 μl spots of these infections were plated to allow estimating how many phages were eluted. Periplasmic extracts were prepared according to standard protocols involving overnight production in 2YT medium and induction with IPTG. Extracts were tested for binding to all 8 Cfa antigens in ELISA. Based on HinfI digestion pattern, amino acid sequences and cross-reactive binding to Cfa antigens, unique candidate genes were selected for further characterization.

### Yeast Library screening

Yeast library screening was performed as previously described (27). For the first round of magnetic-activated cell sorting (MACS), 1×10^10^ of *S. cerevisiae* expressing a surface displayed library of synthetic nanobodies (27) were centrifuged, resuspended in binding buffer (20 mM HEPES pH 7.5, 150 mM NaCl, 2.8 mM CaCl2, 0.05% MNG, 0.005% CHS, 0.1% BSA, 0.2% maltose) and then incubated with anti-fluorescein isothiocyanate (FITC) microbeads (Miltenyi) and FITC labeled MBP for 40 min at 4°C. Yeast were passed through an LD column (Miltenyi) to remove any yeast expressing nanobodies which interacted with the microbeads or MBP. Remaining yeast that flowed through the column were centrifuged, resuspended in binding buffer, and incubated with 1 μM of FITC-labeled N-terminal CfaE protein for 1 h at 4°C. Yeast were then centrifuged, resuspended in binding buffer with anti-FITC microbeads, and incubated for 15 min at 4°C before passing them into an LS column (Miltenyi) and collecting the eluate enriched for CfaE-binding nanobodies. The eluted yeast were expanded and used in a subsequent round of MACS to further enrich for CfaE-binding nanobodies. The second round was performed similarly to the first, but beginning with 4×10^8^ yeast and substituting FITC-labeled CfaE with AlexaFluor647-labeled CfaE, and anti-FITC microbeads with anti-AlexaFluor647 microbeads. High affinity binding yeast were isolated with FACS. In the first round of FACS, yeast binding to 300 nM of AlexaFluor488 labeled CfaE were collected. These yeast clones were grown and subjected to second round of FACS in the presence of human monoclonal antibody (29) that was shown to bind in the proximity of the receptor binding domain. Clones that were outcompeted from binding to the antigen were collected. About 400 individual clones from two rounds of FACS were then grown, stained in a 96-well plate, assessed via flow cytometry for binding specificity to CfaE, and sequenced. 30 clones with unique sequences were isolated and chosen for further characterization, that included further panning with FITC-labeled class 5 adhesins CS1 (class5b) and CS2 (class5c).

### Nanobody purification

Nanobody sequences were cloned into pET-26b vector (EMD Millipore, Cat #69862) to express as a fusion protein with a C-terminal 6xHis tag. Sequence-verified clones were transformed into T7 Express lysY BL21 *E. coli*. Bacteria were grown in Terrific Broth containing 1 mM MgCl2 and 0.01% glucose to an OD600 = 0.7 before induction with 1 mM IPTG. Cells were harvested after an overnight incubation at 27°C. Following osmotic shock, nanobodies were purified from the periplasmic fraction by Ni-NTA chromatography (Gold Biotechnology) and dialyzed against PBS to remove imidazole.

### ETEC strains

H10407 expressing CFA/I fimbriae was purchased from ATCC (ATCC 35401). ETEC strain H10407 was cultured on 2% agar containing 1% Casamino Acids (Sigma) and 0.15% yeast extract (Fisher Bioreagents) plus 0.005% MgSO4 (Sigma) and 0.0005% MnCl2 (Sigma) (CFA agar plates) overnight at 37°C. A total of 1×10^8^ CFU/ml were resuspended in 20% glycerol (Sigma) in phosphate-buffered saline (PBS) solution and kept frozen at −80°C until needed. Strains expressing other adhesin molecules were obtain from University of Maryland. These strains included: # 200145 (CS1), #201546 (CS2), #503046 (CS4), #400599 (CS14), #700056 (CS17), #204648 (CS19), #100483 (CS3), #204348 (CS5), #100001 (CS6) and #100171 (CS21). The growth conditions are the same as H10407.

### Generation of Multimerization and VHH-IgA fusion

Nanobodies were multimerized to dimeric or trimeric forms with (G4S) linkers. The dimeric forms were generated using 6X (G4S) linker to connect two monomeric VHHs in tandem N terminus to C terminus orientation. Trimers were generated using two 3X (G4S) linkers between monomeric nanobody units. The multimers were cloned into pET-26b vector adding a C-terminal 6xHis tag. Nanobodies were purified from the periplasmic fraction by Ni-NTA chromatography (Gold Biotechnology) and dialyzed against PBS to remove imidazole. To generate VHH-IgA fusionbodies, monomeric nanbody sequences were cloned into a pcDNA 3.1 vector containing heavy chain constant region of IgA1 or IgA2 chains without the CH1 domain. Each vector was transformed in NEB5 competent cells, and sequences were verified ahead of transient transfection.

### ELISA

96-well plates (Nunc) were coated overnight at 4°C with 2 μg/ml of purified MBP–CfaE-N. The plates were blocked with 1% BSA plus 0.05% Tween 20 in PBS. Purified nanobodies were diluted in 1 PBS and added to the plates for 1h. The plates were stained with hydroxy peroxidase -conjugated rabbit anti-camelid IgG Fc (1:10,000) for 1 h and developed using TMB Peroxidase substrate (SeraCare). Absorbance at an optical density at 450 nm (OD450) was measured on an Emax precision plate reader (Molecular Devices).

### Flow cytometry

Binding of the yeast surface expressing nanobodies to fluorescently labeled antigens was determined as described previously (27). Briefly, single clone pools were induced by Galactose in Trp-media. 2×10^6^ cells were stained in selection buffer (27) in the presence of anti-AlexaFluor647 labeled HA antibody to monitor clone expression, and FITC labeled antigens with the final concentration of 100 nM. To determine whether nanobody was competing with functional antibody, antibody 68-61 at a concentration of 100 fold of reported Kd was included in the staining mixture. Yeast cells were washed, resuspended in the selection buffer and subjected to flow cytometry on MACSquant.

### Mannose-resistant hemagglutination assay of human group A erythrocytes

H10407 strain cultures were taken from frozen cell banks and diluted in a sterile 0.15 M saline solution until an OD600 of 1 was reached for the assay. Other ETEC strains except #400599 (CS14) were cultured in ETEC medium containing 1% Casamino Acids (Sigma) and 0.15% yeast extract (Fisher Bioreagents) plus 0.005% MgSO4 (Sigma) and 0.0005% MnCl2 (Sigma) and 0.5 mg/ml of Deferoxamine (Sigma Aldrich). CS14-expressing strain (#400599) was cultured on 2% agar containing 1% Casamino Acids (Sigma) and 0.15% yeast extract (Fisher Bioreagents) plus 0.005% MgSO4 (Sigma) and 0.0005% MnCl2 (Sigma) (CFA agar plates) overnight at 37°C plates. Bacteria was resuspended in 1x PBS and dilutions were used to determine the concentration needed to inhibit hemagglutination.

Type A-positive human erythrocytes stored in K3EDTA were washed three times with 0.15 M saline solution and resuspended in the same solution to a final concentration of 1.5% (vol/vol). In a U-bottom 96-well plate (Nunc Thermo Scientific), 100 μl of nanobody was added in duplicate to the top row and diluted 1:2 down the plate in a 0.15 M saline solution. Fifty microliters of appropriately diluted ETEC was added to each well together with 50 μl of a 0.1 MD-mannose solution (Sigma). The plate was incubated for 10 min at room temperature. After incubation, 50 μl of blood solution was added to the plate and mixed well (200 μl final volume). Plates were allowed to sit stagnant at 4°C for 2 h. Hemagglutination was then observed without the aid of magnification. The absence of a pellet of erythrocytes at the bottom of the well is indicative of positive hemagglutination. Blood was ordered fresh every other week (Bioreclamation IVT).

### Caco-2 adhesion assay

Caco-2 cells seeded at 1×10^4^ cells/ml were grown in 96-well tissue culture plates containing Dulbecco’s modified Eagle’s medium (DMEM), at 37°C in 5% CO2 statically. H10407 strain of ETEC was grown overnight at 37°C in ETEC medium containing 1% Casamino Acids (Sigma) and 0.15% yeast extract (Fisher Bioreagents) plus 0.005% MgSO4 (Sigma) and 0.0005% MnCl2 (Sigma). The next day, bacteria were resuspended in PBS and diluted until an OD600 nm of 0.4 was reached. Antibody dilutions were set up in a deep well plate. Antibody dilutions and bacteria were combined at a 1:10 ratio and allowed to shake at 300 rpm for 1 h at room temperature. Meanwhile, Caco-2 cells were washed and incubated in antibiotic free DMEM containing 500 μg/ml of nanobody. After incubation, 0.035 ml of the mixture of antibody and bacteria was added to each well containing Caco-2 cells. The cells were then incubated statically for 3 h at 37°C. The cells were then washed four times with 1 ml PBS to remove non adherent ETEC cells and intensity of luciferase signal was determined.

### Mouse intestine colonization assays

Six-to eight-week-old DBA/2 mice were pretreated with streptomycin (5 g/liter) in the drinking water for 24 to 48 h. Twelve hours prior to bacterial administration, the water bottle was replaced with regular drinking water. One hour prior to bacterial administration, mice received cimetidine (50 mg/kg) intraperitoneally to reduce the effect of stomach acid on ETEC. A total of 1×10^7^ CFU of ETEC strains diluted in PBS were incubated with 100 mg/kg of monomeric anti-CfaE nanobody or an irrelevant nanobody 1 h prior to challenge. In pre-mix model, bacteria and nanobody were mixed and administered in 200 μl volume by oral gavage using 20-gauge bulb-tip feeding needles. In pre-treatment model, mice were pre-treated with nanobody in 200 μl volume in PBS via oral gavage. One or two hours later, bacteria was administered in 100 μl PBS by oral gavage. The mice were allowed to survive for 24 h. At 12 h before euthanasia, food was withdrawn. Following isolation of the small intestine, two segments of ileum (3 cm each), beginning within 0.5 cm of the ileocecal junction and extending proximally 6 cm, were removed and placed in 1 ml sterile PBS (40). Tissues were mechanically homogenized. Samples were serially diluted on MacConkey agar plates and incubated overnight at 37°C. Bacterial CFU were counted the next day.

### Epitope mapping

BioLuminate software (Schrödinger) was used to identify CfaE residues involved in antibody-antigen recognition. A total of 17 amino acids predicted by the software to be involved in the interaction between anti-CfaE nanobodies and the N-terminal portion of CfaE were individually by Genscript. The genes were cloned into pMAL-C5x vector, and the resulting 17 constructs were transformed, expressed, and purified as described above. An ELISA was performed to determine binding of the nanobodies to the mutant proteins in comparison to that to the wild type.

### Statistical analysis

Statistical calculations were performed using the software Prism version 7.03 (GraphPad Software, La Jolla, CA). Comparisons between the hemagglutination or Caco-2 titers of respective antibodies were performed using one-way analysis of variance (ANOVA) with Bonferroni correction for post hoc multiple comparisons.

