## Supplemental Figures and Table for "Oral administration of a single anti-CfaE nanobody provides broadly cross-protective immunity against major pathogenic Enterotoxigenic *Escherichia coli* strains"

**(Supplemental data)**

Alla Amcheslavsky^1^, Aaron Wallace^1^, Monir Ejemel^1^, Qi Li^1^, Conor McMahon^2^, Matteo Stoppato^1^, Serena Giuntini^1^, Zachary A. Schiller^1^, Jessica Pondish^1^, Jacqueline R. Toomey^1^, Ryan Schneider^1^, Jordan Meisinger^1^, Raimond Heukers^3^, Andrew C. Kruse^2^, Elieen M. Barry^4^, Brian Pierce^5^, Mark S. Klempner^1^, Lisa A. Cavacini^1^, Yang Wang^1^

^1^MassBiologics, University of Massachusetts Medical School, Boston, Massachusetts, USA

^2^QVQ B.V. Utrecht, the Netherlands.

^3^Department of Biological Chemistry and Molecular Pharmacology, Blavatnik Institute, Harvard Medical School, Boston, Massachusetts, USA

^4^Center for Vaccine Development, University of Maryland School of Medicine, Baltimore, Maryland

^5^Institute for Bioscience & Biotechnology Research, University of Maryland School of Medicine, Baltimore, Maryland

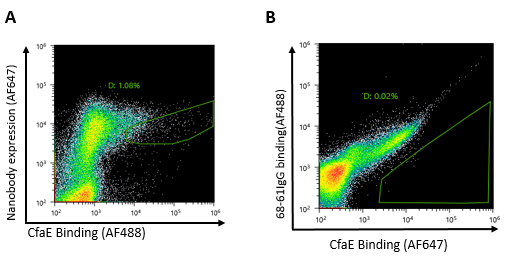

**Supplemental figure 1. Strategy for selection of CfaE binding clones from yeast display library**

(A) High-affinity binders were enriched by fluorescent activated cell sorting (FACS) with

300 nM AlexaFluor488 labeled CfaE. Nanobody expression was monitored with AlexaFluor647 labeled anti-HA antibody.

(B) Competitive FACS screen was performed against a potent anti-CfaE human monoclonal antibody (HuMab), 68-61, to identify nanobodies competing for the binding to putative receptor pocket of CfaE.

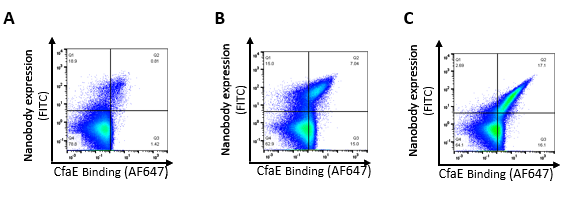

**Supplemental figure 2. Yeast surface displayed clones show various binding affinities to CfaE protein**

Flow analysis demonstrating examples of weak (A), moderate (B) and strong (C) binding of yeast displayed nanobodies to CfaE protein. Single nanobody clones displayed on yeast surface were labled with 100nM of AlexaFluor647 labeled CfaE. Nanobody expression was monitored with FITC labeled anti-HA antibody.

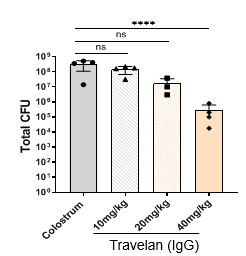

**Supplemental figure 3.** **Effect of Travelan on ETEC colonization in one hour pre-treatment model**

Travelan was used at 10, 20 and 40 mg/kg one hour prior to administration of 10^7^ CFU of H10407. The dose of 40 mg/kg resulted in 3.8 log reduction in colony numbers (*P<*0.0001).

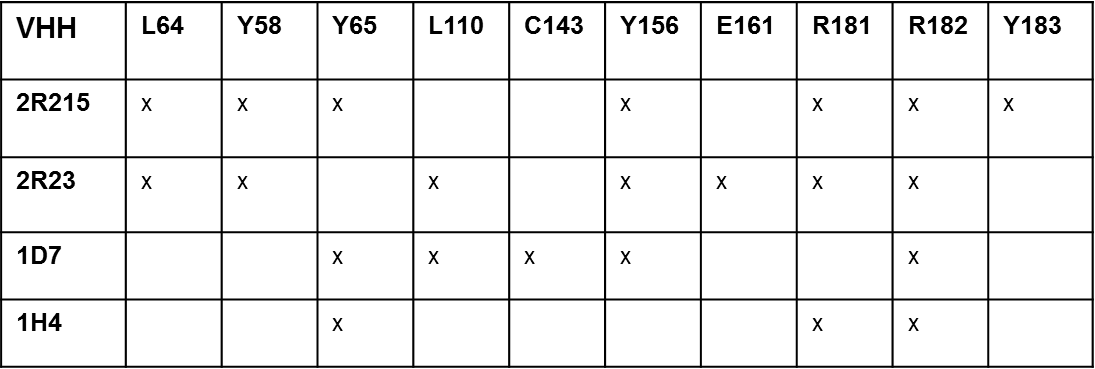

**Supplemental Table 1. Residues required for VHH binding to conserved epitopes on class 5 colonization factor adhesins**

Requirement of residues for VHH binding was examined in ELISA by substituting amino acids to alanine. ELISA results showed that mutating highly conserved residues Y65, R67, E161, D184, R181, R182, Y183, L64, Y58, Y65, L110, Y156 and C143 affected the binding of nanobodies to CfaE.
